# High-performance pipeline for MutMap and QTL-seq

**DOI:** 10.1101/2020.06.28.176586

**Authors:** Yu Sugihara, Lester Young, Hiroki Yaegashi, Satoshi Natsume, Daniel J. Shea, Hiroki Takagi, Helen Booker, Hideki Innan, Ryohei Terauchi, Akira Abe

## Abstract

Bulked segregant analysis implemented in MutMap and QTL-seq is a powerful and efficient method to identify agronomically important loci. However, the previous pipelines were not user-friendly to install and run. Here, we describe new pipelines for MutMap and QTL-seq. These updated pipelines are approximately 5-8 times faster than the previous pipeline, are easier for novice users to use and can be easily installed through bioconda with all dependencies.

## 1 Introduction

Bulked segregant analysis, as implemented in MutMap (Abe *et al.*, 2012) and QTL-seq (Takagi *et al.*, 2013), is a powerful and efficient method to identify agronomically important loci in crop plants. MutMap requires whole-genome resequencing of a single individual from the original cultivar and the pooled sequences of F_2_ progeny from a cross between the original cultivar and mutant. MutMap uses the sequence of the original cultivar to polarize the site frequencies of neighboring markers and identifies loci with an unexpected site frequency, simulating the genotype of F_2_ progeny.

QTL-seq was adapted from MutMap to identify quantitative trait loci. It utilizes sequences pooled from two segregating progeny populations with extreme opposite traits (e.g. resistant vs. susceptible) and single whole-genome resequencing of either of the parental cultivars. While the original QTL-seq algorithm did not assume a highly heterozygous genome, a “modified QTL-seq” has been developed to handle this situation using high resolution mapping (Itoh *et al.*, 2019).

Despite their usefulness, these programs were not user-friendly to install or run and required multiple user inputs. Another problem was that the programs required Coval (Kosugi *et al.*, 2013) for variant calling, which relied on older versions of SAMtools (before 0.1.8).

In this study, we describe newly developed pipelines for MutMap and QTL-seq with updated features.

## 2 Implementation

The new pipelines support read trimming by Trimmomatic (Bolger *et al.*, 2014), replacing fastx-toolkit in the previous pipeline. Trimmed reads are aligned by BWA-MEM (Li and Durbin, 2009), replacing BWA-SAMPE, BWA-ALN and Coval. Improperly paired reads are filtered by SAMtools (Li *et al.*, 2009). Subsequently, a VCF file is generated by the “mpileup” command implemented in BCFtools (Li, 2011). The user can start the analysis from any point in the process, e.g. - from raw FASTQs, trimmed FASTQs, BAM files, or a VCF file. MutPlot and QTL-plot, which are standalone programs, were developed for postprocessing of VCF files. Low-quality variants in a VCF file are filtered out based on mapping quality and strand bias and the actual and expected SNP-indexes calculated based on the AD (allele depth) value of each sample pool (Abe *et al.*, 2012). In QTL-seq, a ΔSNP-index is calculated by subtracting one SNP-index from the other (Takagi *et al.*, 2013). As an option, multiple testing correction (Huang *et al.*, 2019) was also adopted to the simulation. Both pipelines ignore the SNPs which are missing in the parental sample. Candidate causal mutations in the VCF file are shown graphically after executing SnpEff (Cingolani *et al.*, 2012). The procedures are connected by a Python script.

## 3 Results and Conclusions

To compare the performance of the new and old pipelines, we ran MutMap and QTL-seq using four test datasets on an AMD EPYC 7501 processor (Base 2.0 GHz) with 48 GB RAM and 12 threads [located at ROIS National Institute of Genetics in Japan]. The new MutMap and QTL-seq pipelines are approximately 5-8 times faster than the previous pipelines. The ability of the updated pipeline to use a wider range of input file formats reduces the time required for file-management and data handling and makes it easier to use the software.

Greatly reduced processing times for the updated pipelines were accomplished by utilizing more applications with parallel processing (Trimmomatic, SAMtools and BCFtools) and omitting the creation of a consensus FASTA file that had been implemented in the previous pipelines (Fig. 1). Further time-savings were accomplished with the new pipeline by removing user interactions that were required in the previous version. Although the numbers of SNPs plotted were slightly different, the results of the old version and the new version were similar or had slightly better confidence index values (Supplementary figure).

**Fig. 1.**
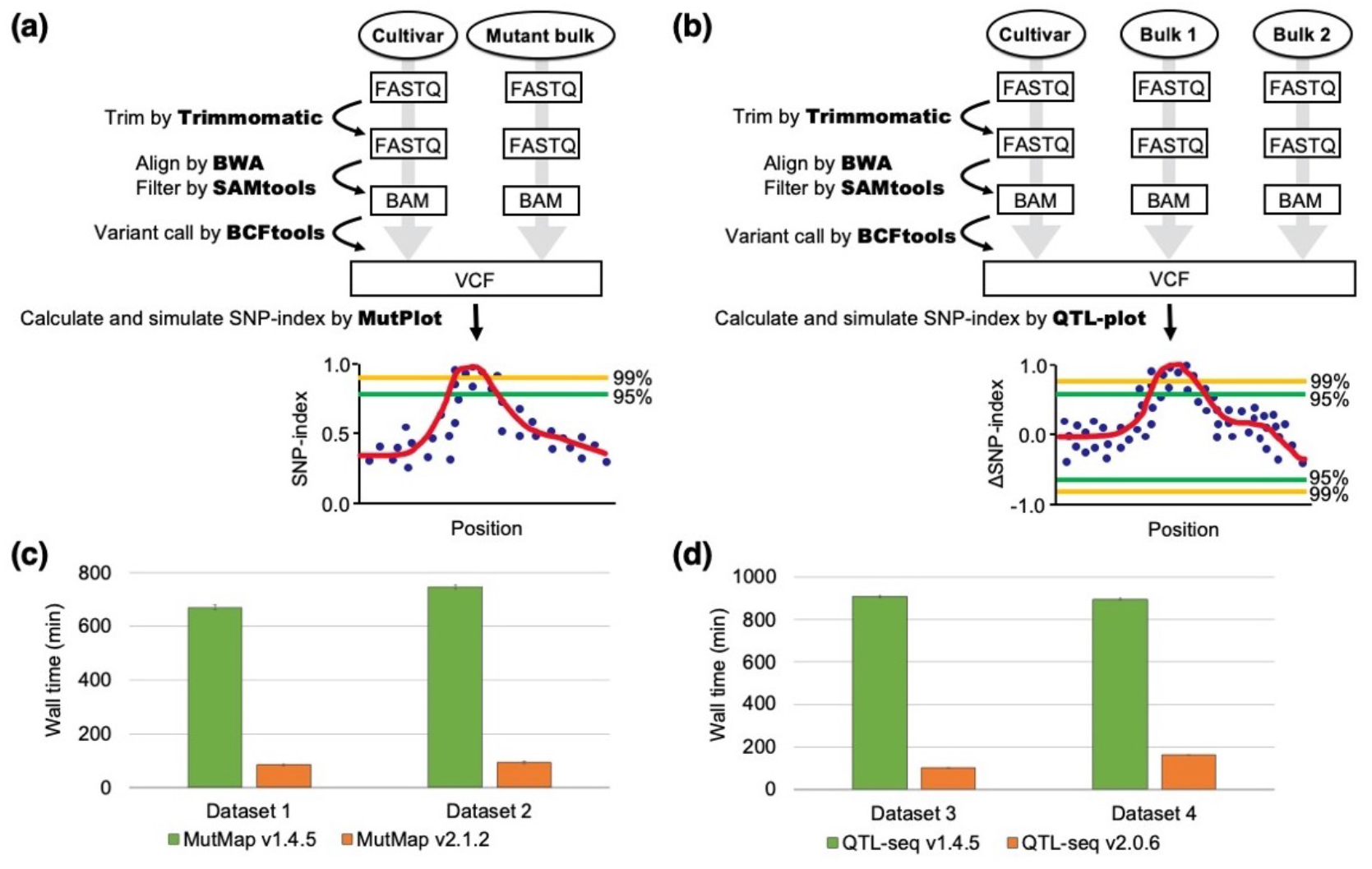
Pipeline workflow and performance of MutMap and QTL-seq. (**a**) The pipeline workflow of MutMap. (**b**) The pipeline workflow of QTL-seq. (**c**) Speed comparison between new (v2.1.2) and old (v1.4.5) pipeline of MutMap. Dataset 1 (Hit1917-pl) and Dataset 2 (Hit1917-sd) can be downloaded as follows: DRR004451; an original rice cultivar “Hitomebore”, DRR001785; mutant bulk of Hit1917-pl, DRR001787; mutant bulk of Hit1917-sd (Abe *et al.*, 2012). (**d**) Speed comparison between new (v2.0.6) and old (v1.4.5) pipeline of QTL-seq. Dataset 3, which was obtained from recombinant inbred lines (RILs) derived from a cross between Nortai and Hitomebore, and Dataset 4, which was obtained from F_2_ progeny derived from a cross between Hitomebore and WRC57, can be downloaded as follows: DRR004451; a parental rice cultivar “Hitomebore”, DRR003237 and DRR003238; two bulks from RILs, DRR003341 and DRR003342; two bulks from F_2_ progeny (Takagi *et al.*, 2013). The tests were performed on AMD EPYC 7501 processor (Base 2.0 GHz) and 48 GB RAM with 12 threads, and the values are means ± SD (n=3).

Currently, these new pipelines can be installed through bioconda with all dependencies. The new pipelines of MutMap and QTL-seq have improved performance and are more user-friendly to install and run, making them very useful for the purpose of genetics studies.

## Acknowledgements

Computations were performed on the NIG supercomputer at ROIS National Institute of Genetics. Additional runs of the QTLseq pipeline using flax resequencing data (unpublished results) were performed using ComputeCanada infrastructure (www.computecanada.ca).

## Supplementary figure

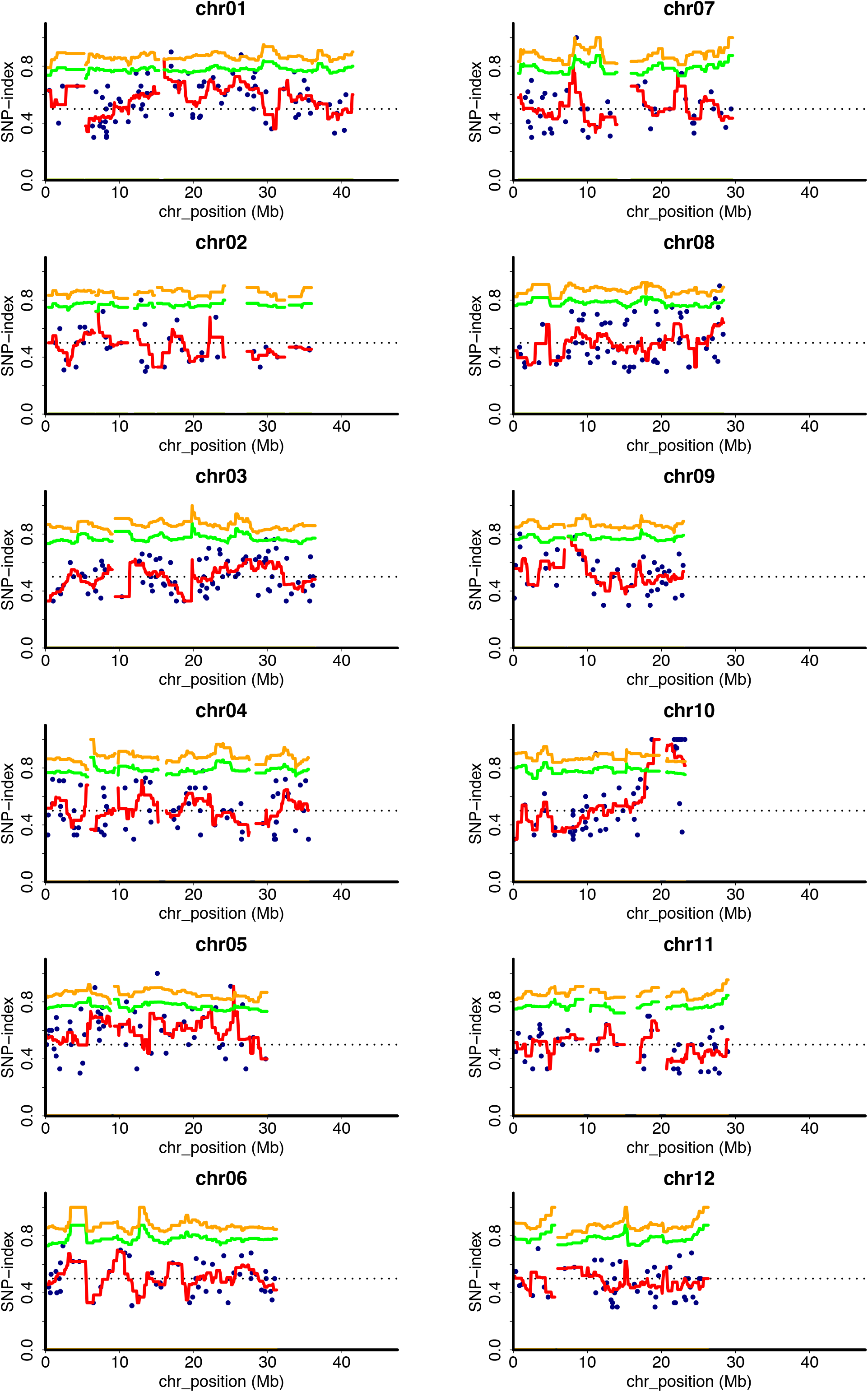
(A) MutMap plot of Hit1917-pl from MutMap v1.4.5 Statistical confidence intervals under the null hypothesis of no QTL are shown (green: *P* < 0.05; yellow: *P* < 0.01).

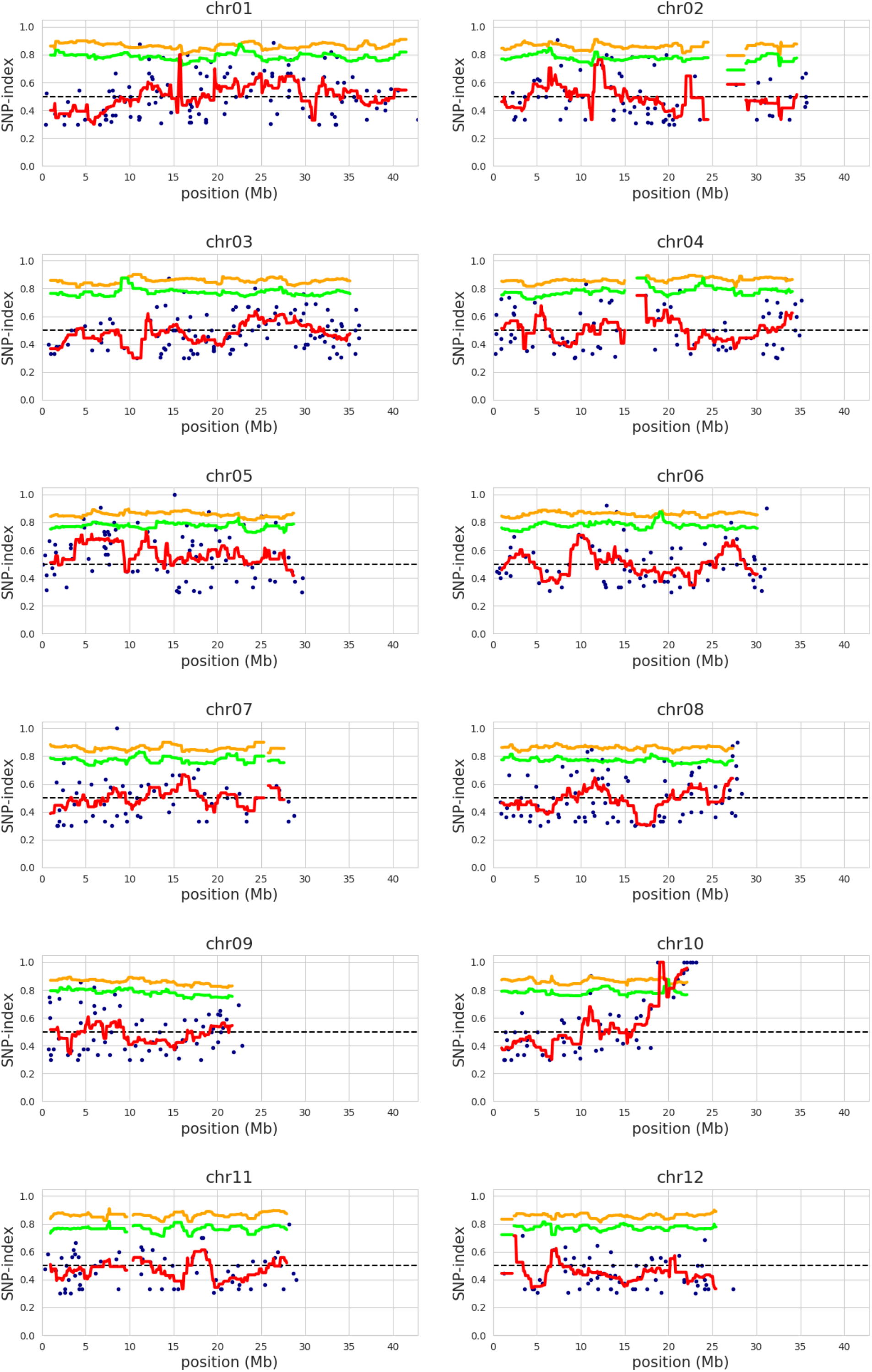
(B) MutMap plot of Hit1917-pl from MutMap v2.1.2 Statistical confidence intervals under the null hypothesis of no QTL are shown (green: *P* < 0.05; yellow: *P* < 0.01).

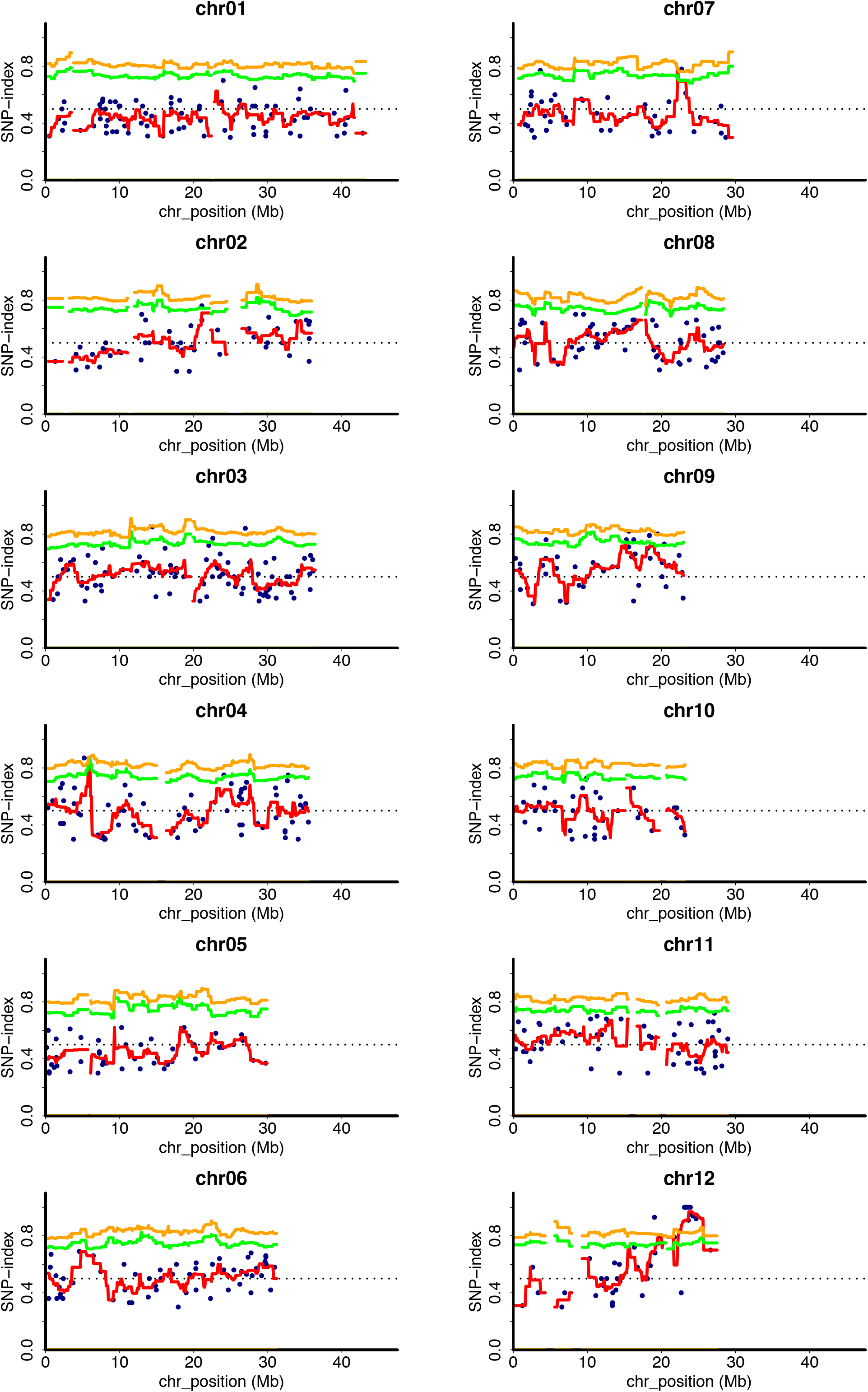
(C) MutMap plot Hit1917-sd from MutMap v1.4.5 Statistical confidence intervals under the null hypothesis of no QTL are shown (green: *P* < 0.05; yellow: *P* < 0.01).

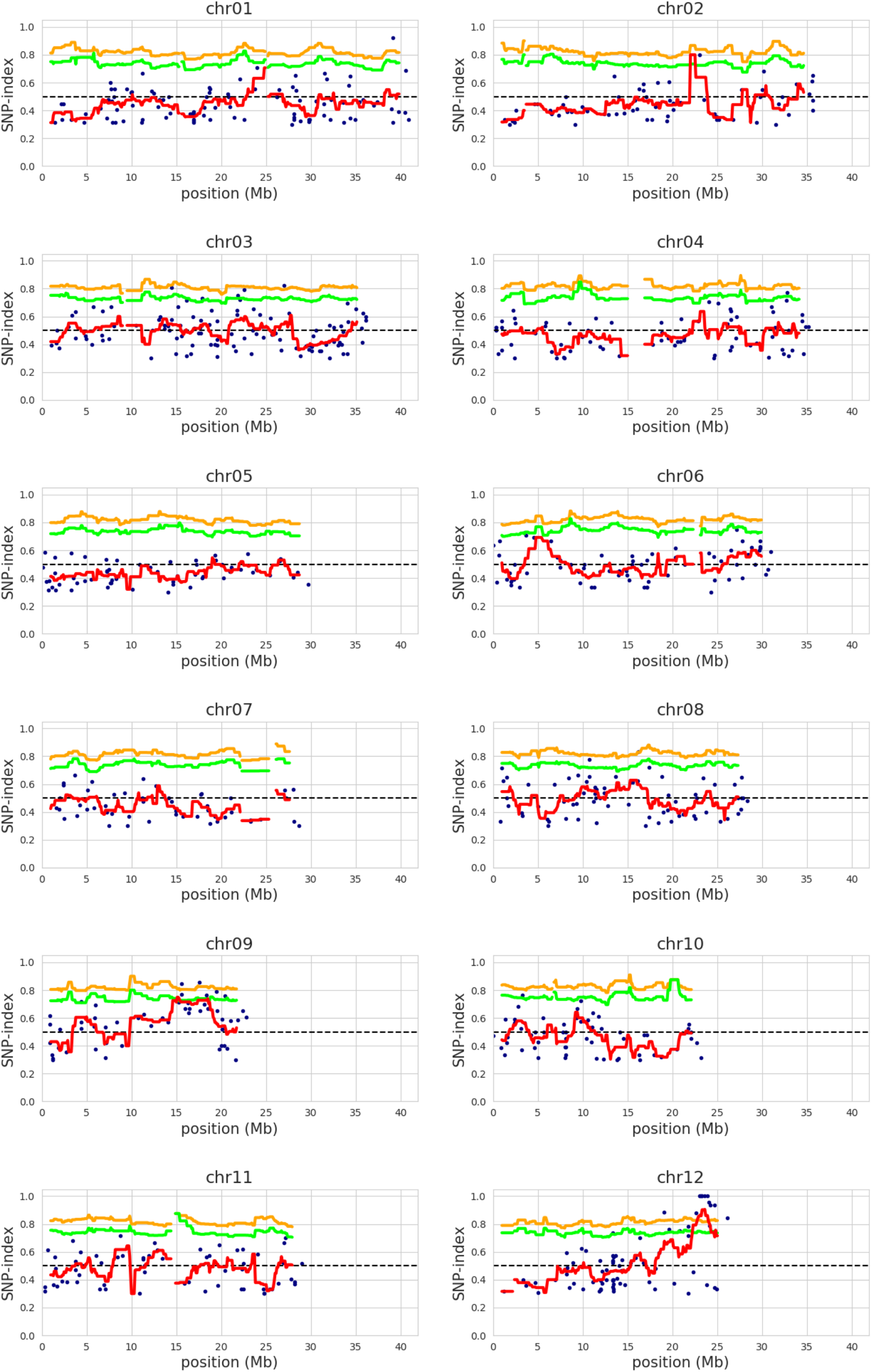
(D) MutMap plot of Hit1917-sd from MutMap v2.1.2 Statistical confidence intervals under the null hypothesis of no QTL are shown (green: *P* < 0.05; yellow: *P* < 0.01).

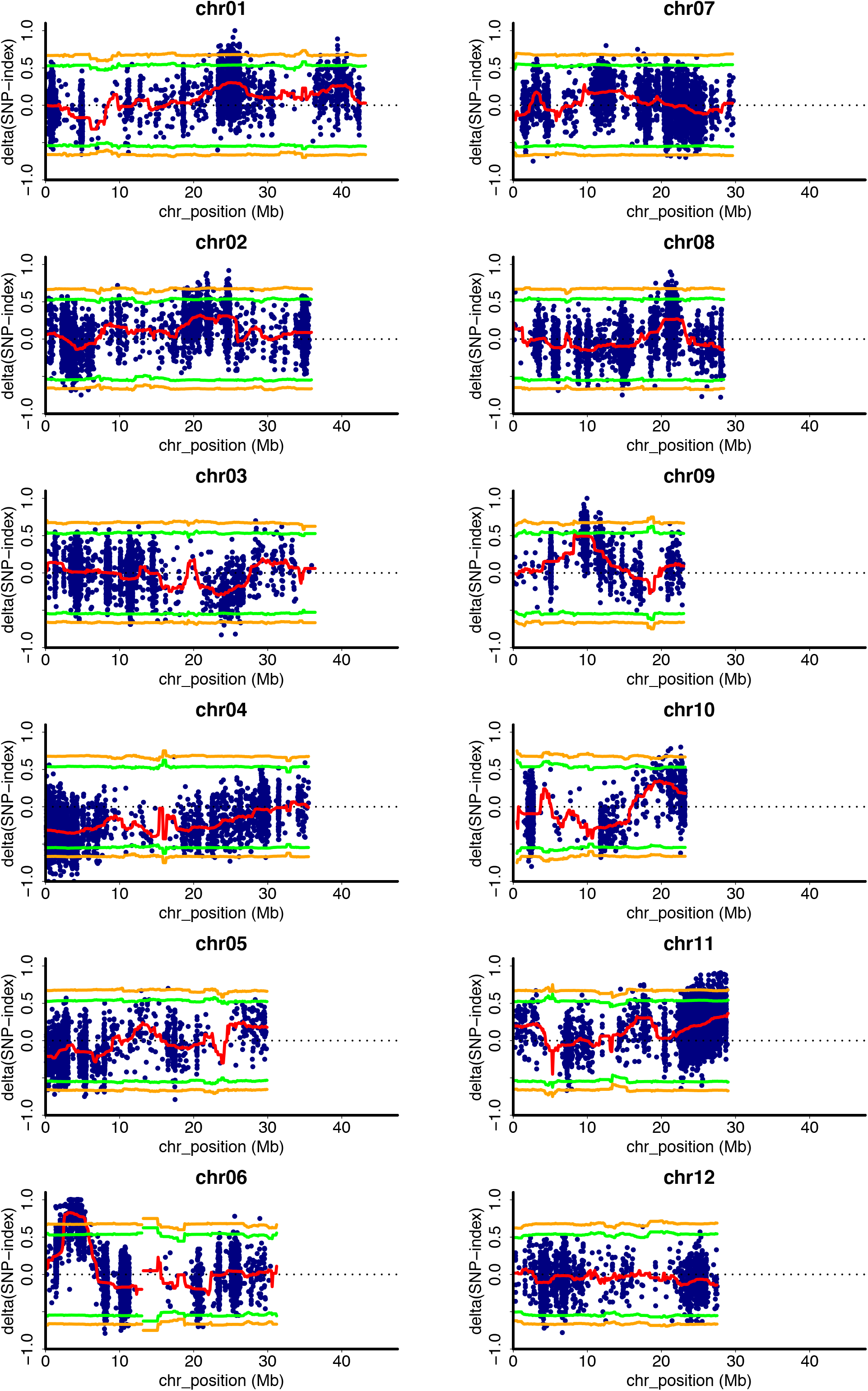
(E) QTL-seq plot of RILs from QTL-seq v1.4.5 The ΔSNP-index plot obtained by subtraction of susceptible-bulk SNP-index from resistance-bulk SNP-index for RILs obtained from a cross between Nortai and Hitomebore. Statistical confidence intervals under the null hypothesis of no QTL are shown (green: *P* < 0.05; yellow: *P* < 0.01).

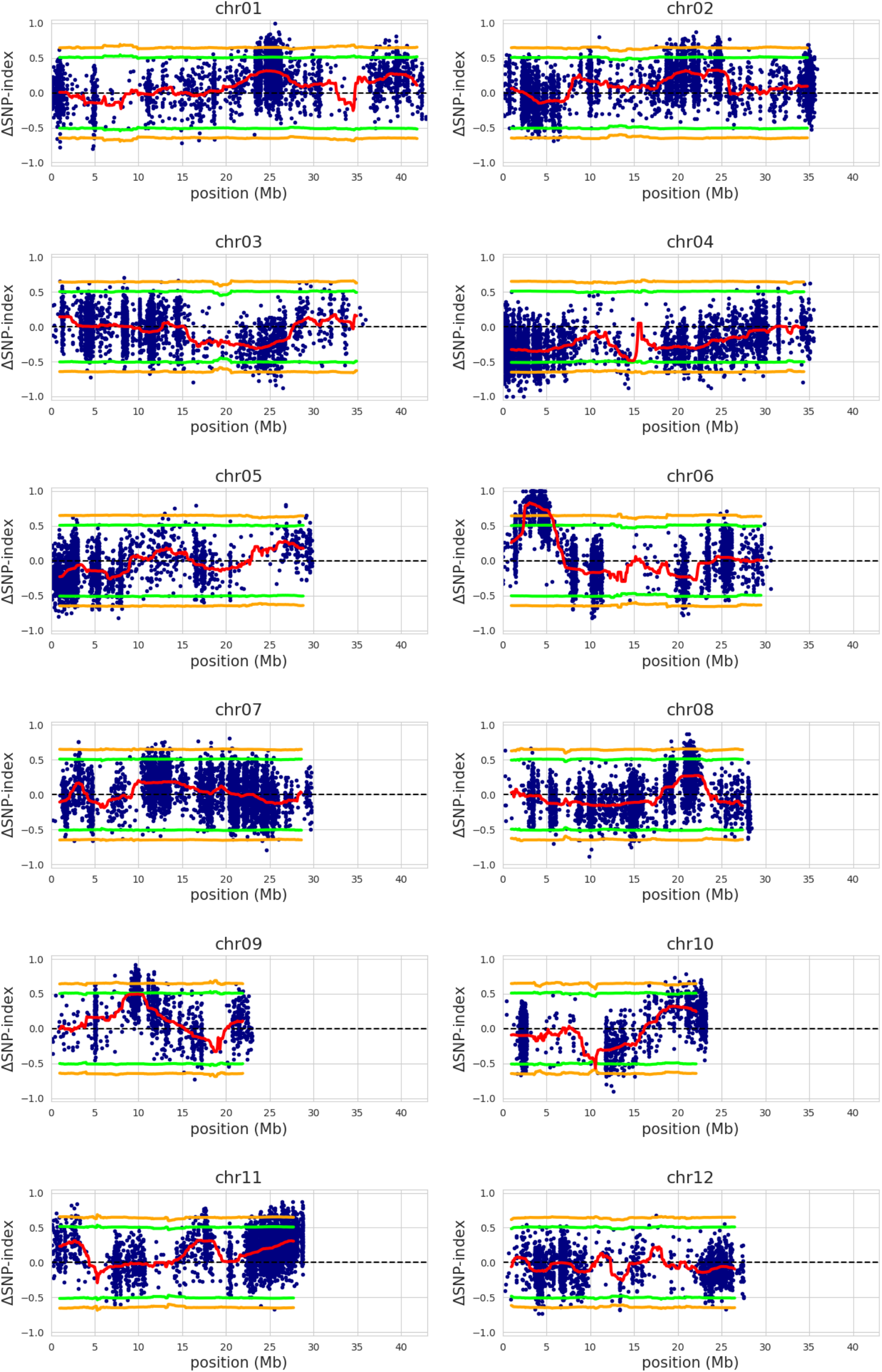
(F) QTL-seq plot of RILs from QTL-seq v2.0.6 The ΔSNP-index plot obtained by subtraction of susceptible-bulk SNP-index from resistance-bulk SNP-index for RILs obtained from a cross between Nortai and Hitomebore. Statistical confidence intervals under the null hypothesis of no QTL are shown (green: *P* < 0.05; yellow: *P* < 0.01).

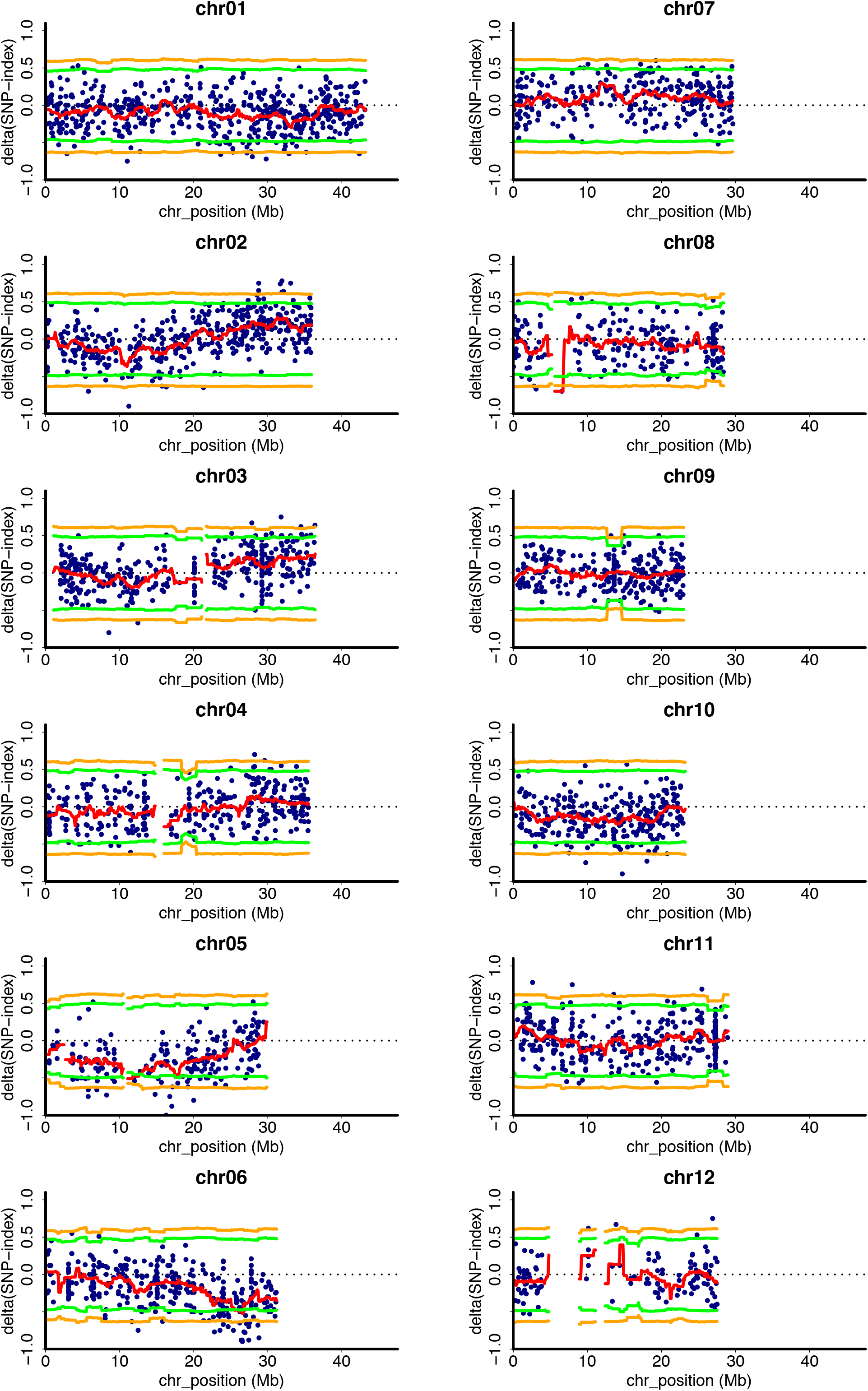
(G) QTL-seq plot of F2 progeny from QTL-seq v1.4.5 The ΔSNP-index plot obtained by subtraction of Highest-bulk SNP-index from Lowest-bulk SNP-index for F2 progeny obtained from a cross between Hitomebore and WRC57. Statistical confidence intervals under the null hypothesis of no QTL are shown (green: *P* < 0.05; yellow: *P* < 0.01).

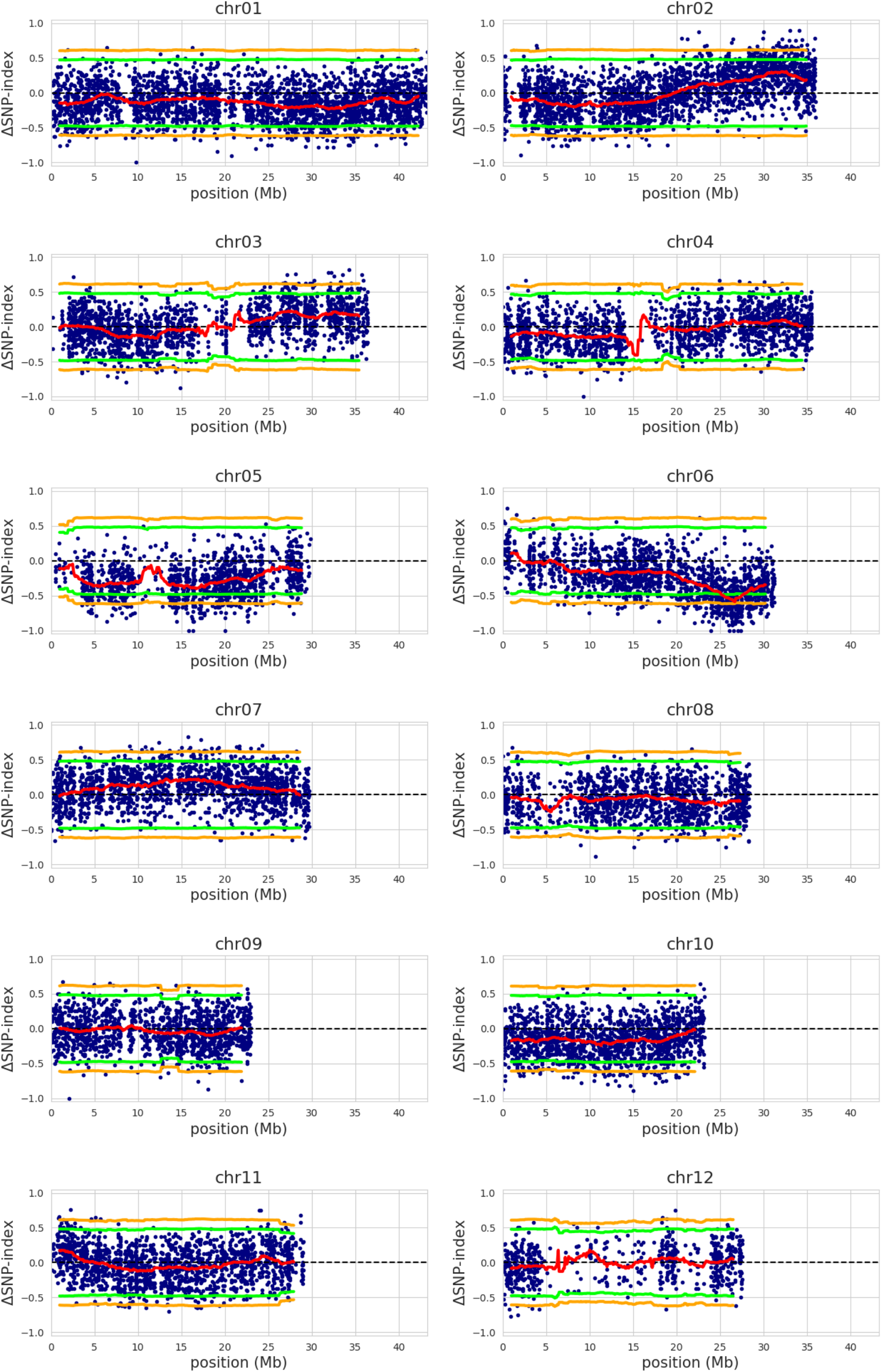
(H) QTL-seq plot of F2 progeny from QTL-seq v2.0.6 The ΔSNP-index plot obtained by subtraction of Highest-bulk SNP-index from Lowest-bulk SNP-index for F2 progeny obtained from a cross between Hitomebore and WRC57. Statistical confidence intervals under the null hypothesis of no QTL are shown (green: *P* < 0.05; yellow: *P* < 0.01).

